# Separating spandrels from phenotypic targets of selection in adaptive molecular evolution

**DOI:** 10.1101/051862

**Authors:** Stevan A. Springer, Michael Manhart, Alexandre V. Morozov

**Affiliations:** Department of Cellular and Molecular Medicine, University of California San Diego, La Jolla 92109, CA, USA; Department of Chemistry and Chemical Biology, Harvard University, Cambridge 02138, MA, USA; Department of Physics and Astronomy, Rutgers University, Piscataway 08854, NJ, USA

**Keywords:** Adaptation, Selection, Pleiotropy, Molecular Evolution, Mutational Accessibility

## Abstract

There are many examples of adaptive molecular evolution in natural populations, but there is no existing method to verify which phenotypic changes were directly targeted by selection. The problem is that correlations between traits make it difficult to distinguish between direct and indirect selection. A phenotype is a direct target of selection when that trait in particular was shaped by selection to better perform a function. An indirect target of selection, also known as an evolutionary spandrel, is a phenotype that changes only because it is correlated with another trait under direct selection. Studies that mutate genes and examine the phenotypic consequences are increasingly common, and these experiments could estimate the mutational accessibility of the phenotypic changes that arise during an instance of adaptive molecular evolution. Under indirect selection, we expect phenotypes to evolve toward states that are more accessible by mutation. Deviation from this null expectation (evolution toward a phenotypic state rarely produced by mutation) would be compelling evidence of adaptation, and could be used to distinguish direct selection from indirect selection on correlated traits. To be practical, this molecular test of adaptation requires phenotypic differences that are caused by changes in a small number of genes. These kinds of genetically simple traits have been observed in many empirical studies of adaptive evolution. Here we describe how to use mutational accessibility to separate spandrels from direct targets of selection and thus verify adaptive hypotheses for phenotypes that evolve by adaptive molecular changes at one or a few genes.

## 1. Inferring the phenotypic target of selection

### The problems of pleiotropy, correlated traits, and indirect selection

Adaptations are phenotypes shaped by selection to perform a function. But we cannot assume that a trait evolved for the function that we happen to assign it (Williams 1966; Gould and Lewontin 1979; Nielsen 2009). To understand the reason a trait evolved, we must formulate and test adaptive hypotheses – scenarios that specify exactly which phenotypic differences created the fitness differences that drove evolution (Williams 1966; Mayr 1983). Tests of adaptive hypotheses are confounded by two problems. The first is that individual mutations typically affect many traits simultaneously, a phenomenon known as pleiotropy. The second is that mutational effects on different traits can be correlated, and if so, a trait can change toward a particular state even when it has no direct consequence for fitness. Traits can evolve solely because they are correlated with some other trait that is important to fitness. We refer to this apparent selection resulting from correlations between traits as indirect selection, and to traits that evolve in this manner as evolutionary spandrels. To verify an adaptive hypothesis, one must be able to distinguish between a spandrel and a trait that was truly a direct target of selection (Pearson 1903; Lande and Arnold 1983).

If we could empirically determine the spectrum of phenotypes available by mutation, we could make two straightforward predictions: *i*) Traits that are not themselves under direct selection should tend toward phenotypic states that are reached frequently by mutation (Stadler et al. 2001). In other words, in the absence of direct selection, traits should evolve toward phenotypes that are mutationally accessible. If the trait in question is weakly coupled to another trait under direct selection, evolution is expected to move its phenotypes toward more accessible states (although, depending on the strength of correlation with the trait under selection, maximally accessible states may never be reached). *ii*) On the other hand, direct selection pushes traits along evolutionary paths that increase fitness, and can fix beneficial mutations even if they are not easily accessible by mutation alone. Similarly, if a trait persists after a series of mutations, even though most potential mutations change it, then it must be under stabilizing selection. These predictions form the basis of the accessibility test described here – a method that uses mutational accessibility to verify the phenotypic targets of adaptive molecular evolution.

As an example, consider a protein that binds to a ligand but must be correctly folded to do so. The protein has two relevant traits: folding stability (quantified by the fraction of proteins in the cell that are properly folded), and binding strength (quantified by the fraction of proteins in the cell that are bound to a ligand). These two traits are inevitably correlated because a protein can only bind when properly folded, so that the fraction of folded proteins is always greater than or equal to the bound fraction (Fig. 1). In principle, selection could act directly on one of these traits and not the other - solely to improve either the binding interaction, or the stability of folding. But even if selection directly acts on only one of these traits, indirect selection can drive change in the other trait if the effects of mutations on the two traits are correlated (Manhart and Morozov 2015). For example, even if there is selection only for folding (Fig. 1, blue horizontal arrow), the fraction of bound proteins will tend to increase simply because proteins that bind properly are more abundant among proteins that fold properly. In other words, proteins that bind well become more accessible by mutation as folding improves. Improved binding could thus be a spandrel that evolves in the absence of direct selection.

**Figure 1.**
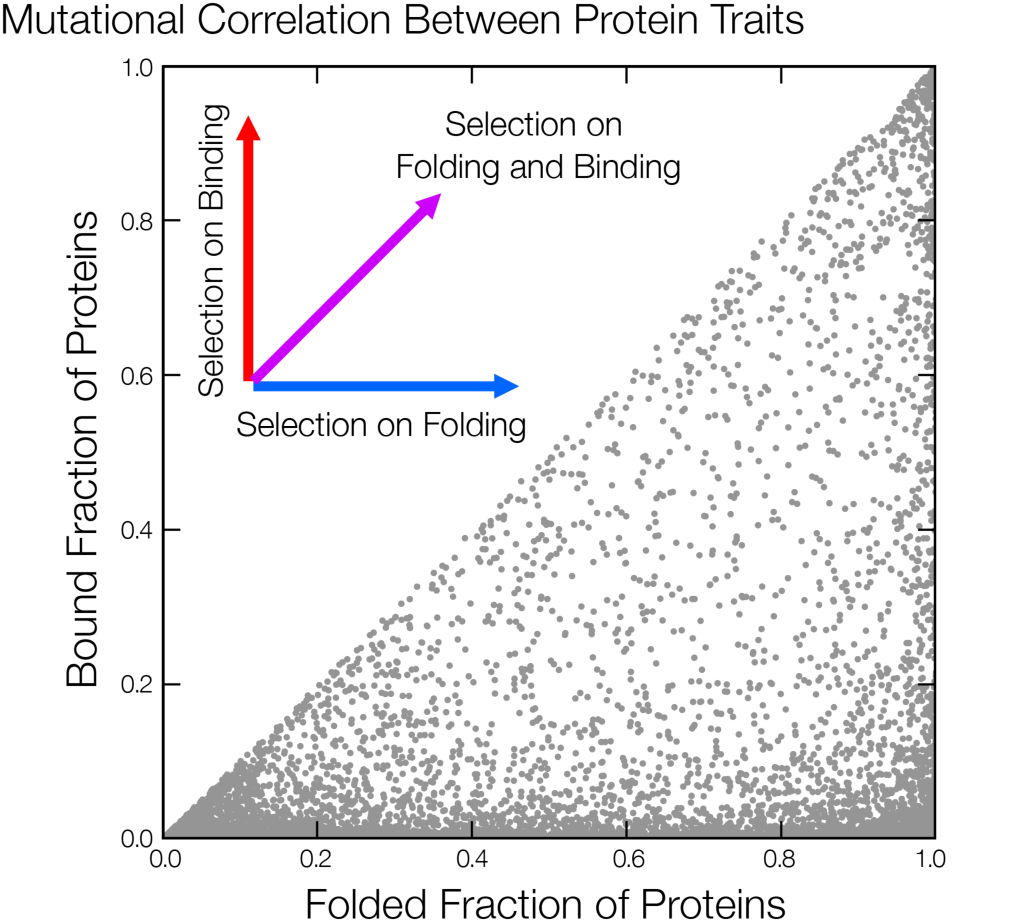
Mutational correlations and indirect selection on protein traits. Distribution of two possible protein traits - folding (quantified by the fraction of proteins that are folded) and binding (quantified by the fraction of proteins that are bound to a ligand) - in a simple thermodynamic model of protein kinetics (Manhart and Morozov 2015). Each residue makes an independent, additive contribution to the free energies of folding and binding (Wells 1990), which in turn determine the folding and binding probabilities via Boltzmann factors. Here we consider protein sequences varying only at 6 sites (e.g., at the binding interface), with a reduced alphabet of 5 amino acid types. The free energy contribution of each amino acid type at each site is randomly sampled from a Gaussian distribution with mean and standard deviation of 1 kcal/mol, consistent with observed distributions of mutational effects (Thorn and Bogan 2001; Tokuriki et al. 2007); total free energies are offset such that mean (over all possible sequences at the binding interface) free energy of folding is 2 kcal/mol and the mean free energy of binding is 5 kcal/mol. Each grey point represents the folding and binding traits of a different protein sequence. These two traits must be correlated because the fraction of folded proteins is always greater than, or equal to, the fraction of bound proteins (since the protein can only bind when properly folded). Arrows indicate different direct selection scenarios: direct selection on binding only (red arrow), direct selection on folding only (blue arrow), and direct selection on both binding and folding (magenta arrow).

Molecular tests of phenotypic accessibility are only useful if we can quantify the space of possible phenotypes available by mutation for a particular trait. For quantitative traits, this difficult problem can be avoided by assuming that any small phenotypic change can be achieved by mutation (Maynard Smith 1978; Grafen 1984) (Fig. 2). Measures of the variances and co-variances of quantitative traits can identify the phenotypic targets of single selective events and find Pareto-optimal compromises between different selective regimes (Lande and Arnold 1983; Crespi 1990; Shoval et al. 2012). However, extending these kinds of analyses from quantitative traits to traits with a simple genetic basis (phenotypic differences caused by mutations at a few specific genomic loci) requires information or assumptions about mutational constraints on phenotypic change.

**Figure 2.**
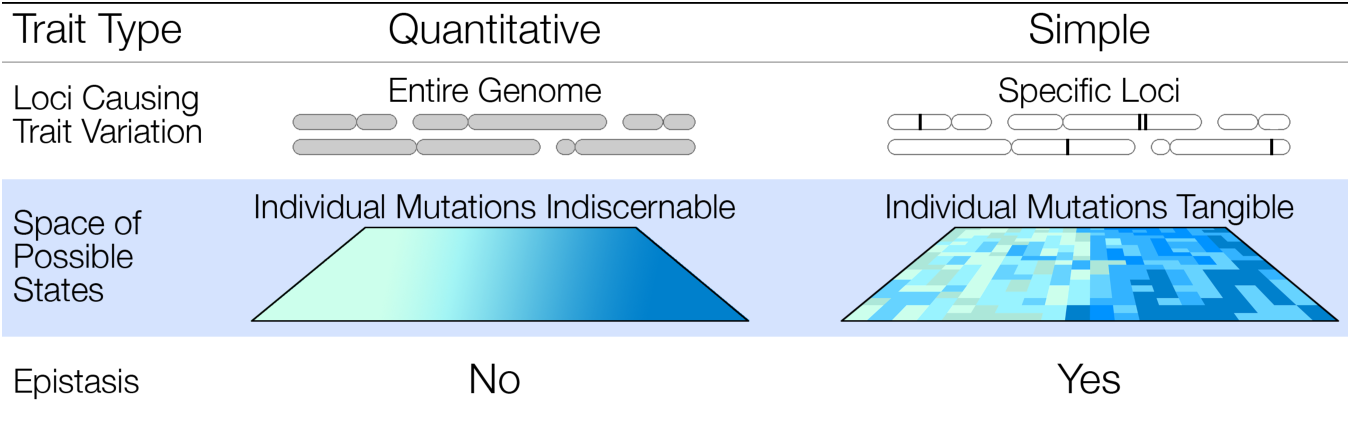
The phenotype spaces of quantitative and simple traits. Finding the phenotypic targets of selection in quantitative traits and genetically simple traits requires different assumptions about phenotype space. Classical optimality models reduce the complexity of phenotype space by assuming that any small phenotypic change can occur, that there is no epistasis, and that the structure of the phenotype space does not change from one mutation to the next. In contrast, the mutational accessibility test focuses on simple traits (traits dependent on just a few genomic loci). In simple traits the size of the phenotype space is reduced, allowing us to study variants of just those loci that cause an observed phenotypic difference. In simple traits, the effects of individual mutations may be epistatic and can significantly alter the space of alternatives available to evolution. Therefore, these traits will benefit most from an explicit genetic investigation.

### Natural adaptations can have simple genetics

A striking empirical finding is that some instances of natural adaptive evolution have a simple genetic basis (Orr and Coyne 1992; Bell 2009; Conte et al. 2012; Martin and Orgogozo 2013; Gallant et al. 2014; Rosenblum and Parent 2014). Natural phenotypic changes sometimes occur by mutations in one or a few genes (Nachman et al. 2003; Bradshaw and Schemske 2003; Hoekstra et al. 2006; Storz et al. 2007), and the same genes or even the same mutations can evolve in parallel (Wichman et al. 1999; Holder and Bull 2001; Cresko et al. 2004; Colosimo et al. 2005; Zhang 2006; Musset et al. 2007; McDonald et al. 2009; Jiang et al. 2012; Shen et al. 2012; Frankel et al. 2012; Springer et al. 2014; Wessinger and Rausher 2015). Artificial mutants of lab organisms can have phenotypes that resemble related species, sometimes due to the same genes or pathways that cause the natural difference (Koufopanou and Bell 1991; Parichy and Johnson 2001; Shapiro et al. 2004; Owen and Bradshaw 2011). Mimics are known to use the same genes as the organisms they model to achieve similar forms, and even regulatory elements can evolve in parallel (Reed et al. 2011; Gallant et al. 2014). Co-evolving proteins maintain their partners over long time intervals (Clark et al. 2009; Hellberg et al. 2012), and conflicts between the same proteins can arise independently in response to similar ecological interactions (Feldman et al. 2012; Dobler et al. 2012; Zhen et al. 2012). Evolution sometimes uses the same genetic elements repeatedly, so for some adaptations there must be only a few loci whose mutations can create an appropriate phenotypic change (Fig. 2). In such cases, it may be possible to move beyond just identifying the genes that cause phenotypic differences, by quantitatively estimating which phenotypic changes are common and which are rare when these causal loci are experimentally mutated. For natural phenotypic differences with simple genetic causes we can thus empirically evaluate the accessibility of the derived phenotype by mutagenesis.

### Distinguishing traits

There are fundamental limitations on our ability to resolve direct and indirect selection. It will not be possible to distinguish the phenotypic targets of selection from their pleiotropic effects when these components of phenotypic variation are strongly correlated (when for any initial state, the maximum phenotypic change in one trait nearly always corresponds to the maximum phenotypic change in another). But selection is also powerless to make this distinction – in terms of evolutionary response, perfectly-correlated phenotypes can in fact be considered one trait (Lewontin 1978; Stadler et al. 2001; Wagner and Zhang 2011). All tests of adaptation (whether quantitative genetic or molecular) thus rest on the fact that we can only separate direct and indirect selection when mutational effects on the measured traits are not too strongly correlated. In other words, for a set of mutations with the same effect on the trait under selection, each must have an independent suite of effects on other traits (Lewontin 1978; Stadler et al. 2001). Fortunately, the extent of correlations between traits can be verified experimentally. For example, directed protein evolution experiments commonly observe weak or no correlations between various biophysical traits: mutations that modify protein-ligand affinity or catalytic ability typically have varying effects on stability, and non-pleiotropic mutations which affect one trait and not another are nearly always available (Bloom and Arnold 2009). Lack of correlation between traits is also observed in natural contexts. For example, mutations of the Agouti locus have a suite of effects on light and dark coloration in mice, but selection can separate one trait from another (Linnen et al. 2013). In this paper, we will assume that when we refer to a trait, we mean the feature being measured, together with all the features that are strongly correlated with it (i.e., with the linear correlation coefficient close to ±1.0). Therefore, strictly speaking, a trait is a set of phenotypic features with correlations that cannot be broken by mutation.

## 2. Practical exploration of phenotype space

For traits with a simple genetic basis, it is possible to recreate the set of mutations that caused a natural phenotypic difference, and explore their phenotypic effects one-by-one (Dean and Thornton 2007; Harms and Thornton 2013). In one of the first examples, Weinreich et al. recreated mutational paths during the evolution of the TEM antibiotic resistance gene (Weinreich et al. 2006). Each path is a different way of ordering the set of mutations that separate the ancestral and derived alleles. The researchers measured each mutation’s phenotypic effect on cefotaxime resistance, and estimated its fixation probability by assuming that selection acted only on this measured trait. Exploring phenotype space in this way is ambitious even for the simplest traits. As the number of differences between the ancestral and derived alleles increases, the number of possible intermediate alleles goes up exponentially, and the number of possible mutational paths increases factorially (Weinreich et al. 2005). To estimate accessibility, we need methods that generate vast molecular diversity and measure its phenotypic consequences. Equally importantly, we need evolutionary models that describe tractable representations of phenotype space.

### Creating and phenotyping molecular diversity with combinatorial molecular biology

Modern molecular biology methods can be used to measure the phenotypic effects of many protein variants, and link each phenotype to its DNA sequence (Scott and Smith 1990). Combinatorial mutagenesis can generate mutational diversity exceeding 10^13^ variants (Weiss et al. 2000; Overstreet et al. 2012). Together, these methods have been used to explore huge spaces of molecular variation, seeking new drug-binding partners, and mapping functional residues in protein-protein interactions. For example, the binding energetics of human growth hormone (hGH) and its receptor (hGHR) have been completely mapped by quantitative saturation mutagenesis: every possible single amino acid mutation of the hGH binding site has been created and tested for receptor affinity (Pál et al. 2006). High-throughput combinatorial methods can recreate ancestral proteins (Zhu et al. 2005; Lunzer et al. 2005; Szendro et al. 2013), find mutations that interconvert the activity of related but functionally-divergent proteins (O’Maille et al. 2008), and explore the function of intermediates between the ancestral and derived state (Weinreich et al. 2006; Bridgham et al. 2009; Field and Matz 2010). Libraries of cis-regulatory regions can measure the effect of regulatory sequence variation on transcription factor binding and protein expression (Gertz et al. 2009). Cell sorters can quickly measure many thousands of cells for any phenotype that can be coupled to a fluorescent marker. Thus practical tools for creating and phenotyping molecular diversity on a large scale exist. To verify which phenotypes were targets of selection, we must use these tools to explore phenotype space in ways that inform us about the accessibility of a phenotypic change (Fig. 3).

**Figure 3.**
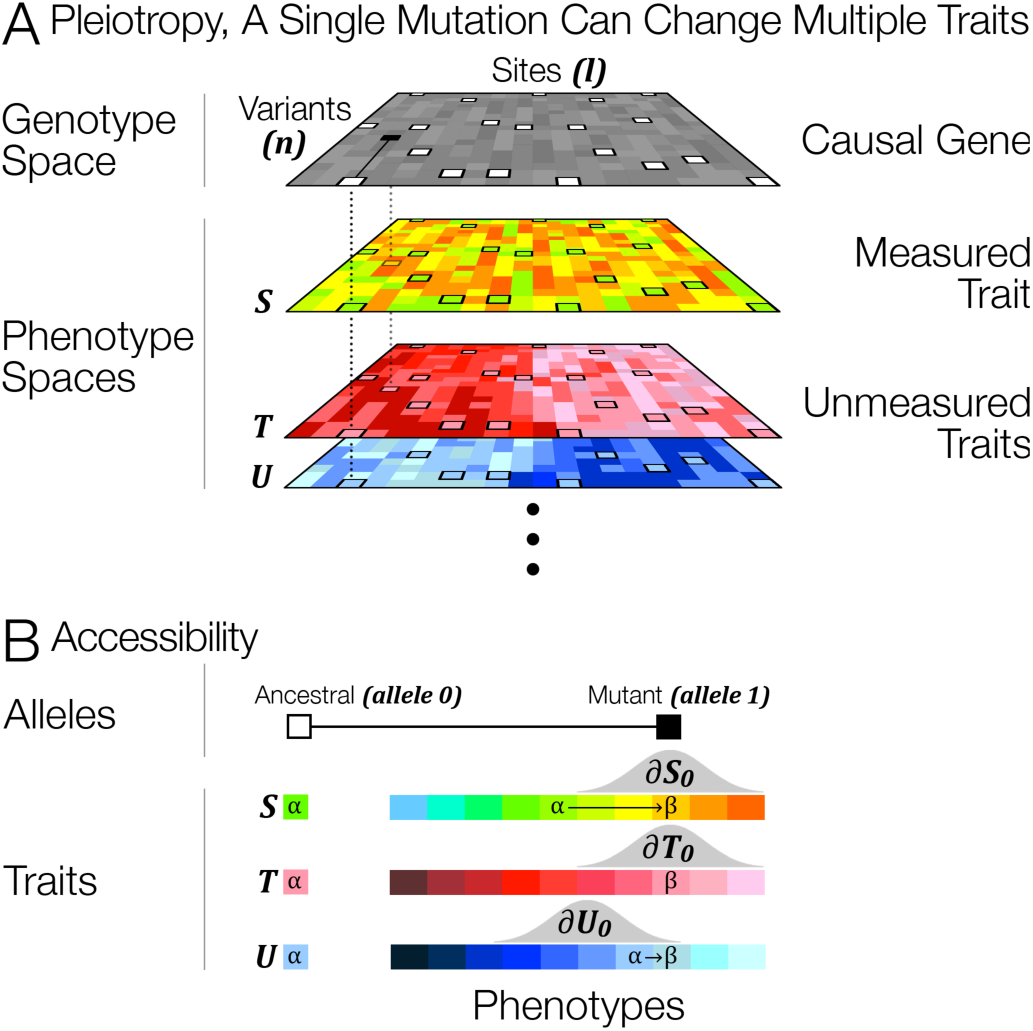
Pleiotropy, accessibility, and evolution by indirect selection. (A) A genotype space and its corresponding phenotype spaces. Each of the ***l*** sites has ***n*** mutational variants. Thus, the mutational neighborhood size (i.e., the number of single-point mutations available to a sequence) is ***l* × *n***, and the total number of sequences is ***n*^*l*^**. For simplicity, we do not consider strongly deleterious mutations, which would result in null phenotypes. White boxes show the wild-type sequence; the black box shows a mutation, which will change the measured phenotype **S**, as well as unmeasured phenotypes ***T*** and ***U***. It is difficult to verify the target of selection because a single mutation can have pleiotropic effects on multiple traits. (B) Mutational neighborhoods of individual alleles are denoted as ∂***Trait*_*allele*_**. Grey frequency distributions represent accessibility: they show how often a particular phenotypic change occurs by a single mutation of an ancestral allele (***allele 0***) for different traits (***S, T, U***). Trait ***S***: most mutations shift the ancestral phenotype green (α) toward orange (β); the observed phenotypic change (from green to orange, α→β) would occur frequently even in the absence of direct selection. Trait ***T*** is neutral with respect to this mutation; its phenotype did not change, nor was it expected to, given that mutations with phenotype α=β are common in ∂***T*_*0*_**. Trait ***U*** evolved in an unexpected direction, away from mutationally accessible states. Selection on this trait may have caused the change observed in trait ***S*** because of their shared genetic basis.

### Tractable representations of phenotype space

To create the most general model of phenotypic evolution, we would need to study every phenotype created by every mutation of the genome, and then consider evolutionary trajectories in this vast phenotypic space (Maynard Smith 1978). This is clearly impossible and will remain so – the number of possible phenotypes and evolutionary trajectories is simply too large to be ever explored completely. If we hope to gain an empirical measure of accessibility, we must reduce the combinatorial complexity and the size of phenotype space. To reduce the number of evolutionary trajectories to be considered, we often assume that selection eliminates or fixes mutations more quickly than the waiting time between mutations (the strong selection, weak mutation, or SSWM, regime). Therefore mutations fix one at a time, and evolution works on the set of phenotypes made available by single mutations of each successively fixed allele (Gillespie 1984; Weinreich et al. 2005). Thus we can explore the neighborhood of single mutations to estimate the null probability of a particular phenotypic change (note that this null is shaped by correlations between traits, but we do not need to explicitly specify the correlated traits). The size of this neighborhood depends on the number of mutations in the genome that influence our trait of interest. Are mutations of small phenotypic effect spread out evenly, as is typically assumed in quantitative trait genetics, or are there a few loci that influence a trait much more than others (Fig. 2)? The approach described in this paper is only needed when genetic constraints significantly influence the course of phenotypic evolution, which is more likely when traits have simple genetic causes (Grafen 1984; Springer et al. 2011). In cases with simple genetics, we can focus on coding and regulatory sequences of the genes known (or suspected) to form the genetic basis of the trait of interest. In the opposite limit of numerous mutations with small phenotypic effects, existing methods from quantitative genetics are capable of distinguishing direct and indirect selection (Lande and Arnold 1983; Shoval et al. 2012).

## 3. The mutational accessibility test

To illustrate the accessibility test, we will use one of the many examples of positive selection whose phenotypic effects can be determined but whose exact evolutionary cause is unknown (Stolz et al. 2003; Nielsen 2009). The ventral and dorsal light organs of Jamaican click beetles (*Pyrophorus plagiophthalamus*) fluoresce in a variety of colors because of variation in their luciferase proteins (Wood et al. 1989). The inferred ancestral luciferase emits green light. Alleles encoding yellow and orange fluorescence have evolved by positive selection, as confirmed by both McDonald-Kreitman and dN/dS tests (Stolz et al. 2003). But what was the phenotypic target that caused this adaptive molecular evolution? Is emitting orange an adaptation, in the sense that alleles producing more orange light conferred higher fitness? Or is the color shift inevitable, a indirect consequence of selection on an unmeasured trait that evolved because most mutations of the luciferase protein change the phenotype toward orange?

We can answer this question by determining the probability of evolving toward orange in the absence of direct selection. If most mutations in the neighborhood of the ancestral click beetle luciferase cause it to glow orange, we are not forced to invoke direct selection on color to explain the switch to orange, as it could have evolved with or without direct selection. However, if these mutations are non-neutral, high mutational accessibility of orange implies that this phenotypic change could have been driven by indirect selection. Selection may have targeted on another trait encoded by luciferase – for example, anti-oxidant activity or protease sensitivity (Thompson et al. 1997; Barros and Bechara 2000). Alternatively, the switch to orange could be both beneficial and highly accessible.

### Single-mutation scenario

If there were only a single mutation separating ancestral and derived alleles, the test would be conceptually simple. Specifically, we could sample a neighborhood of possible mutations of the ancestral sequence by mutagenesis. When expressed in E. coli, luciferase proteins fluoresce at the same wavelength as their natural equivalents (Stolz et al. 2003). We can thus measure the effects of individual mutations by cloning a library of artificial mutations *en masse* into individual E. coli cells, and phenotyping each variant with a fluorescence activated cell sorter (FACS). The aim is to compare the phenotype of the natural mutation with the phenotypes in the neighborhood of possible single mutations to determine if the natural mutation is atypical. Specifically, we could use the neighborhood to assign a p-value to the probability of a random mutation occurring with a phenotypic change at least as extreme as the one observed. A low p-value would thus imply that a mutation with the observed effect occurs rarely in the absence of direct selection. Empirical maps of mutational neighborhoods allow us to estimate accessibility (i.e., the probability of moving from one phenotypic state to another) in our null model of evolution by indirect selection, and therefore to assess whether or not we need to invoke direct selection to explain an observed phenotypic difference (Fig. 4).

**Figure 4.**
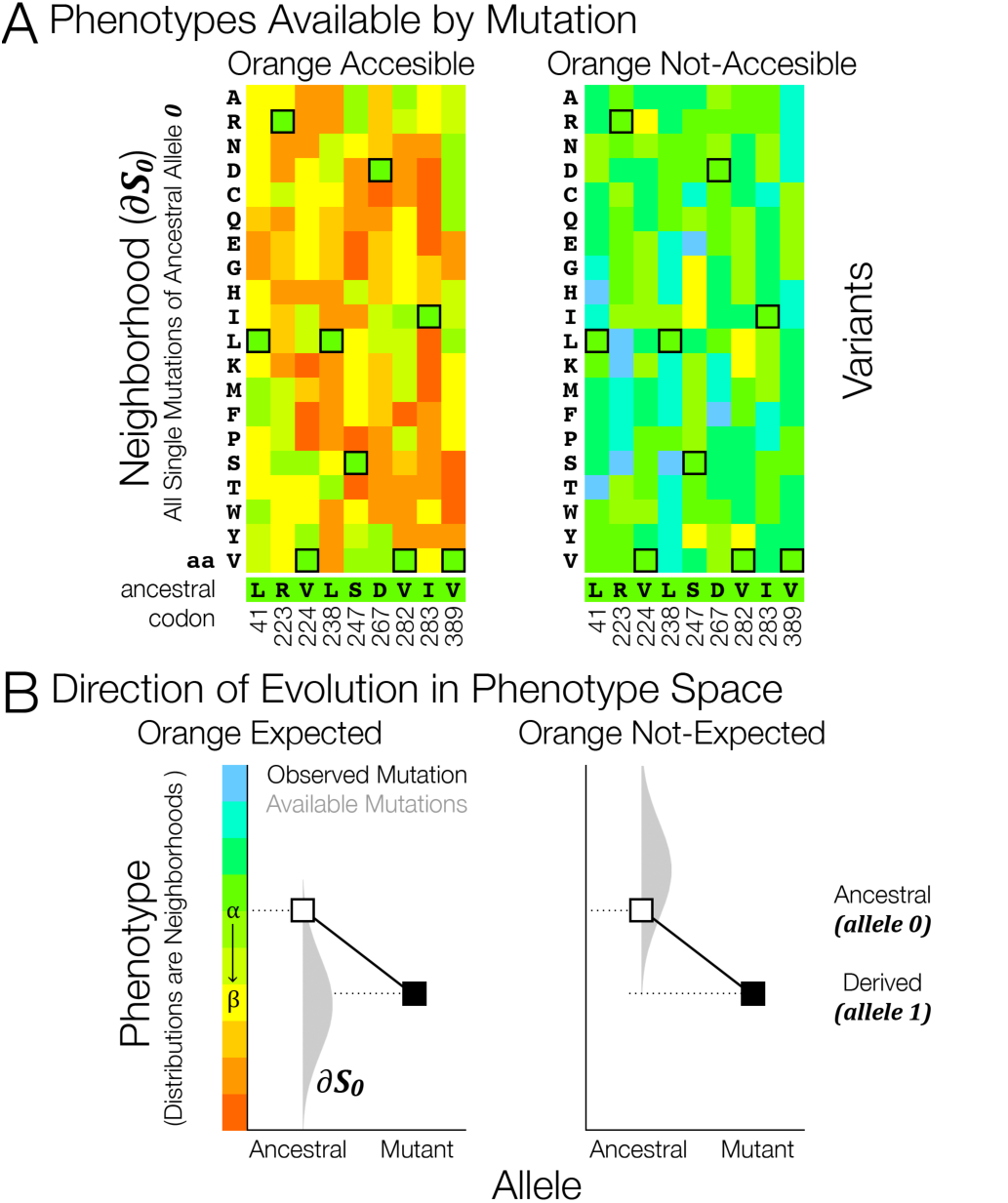
Distribution of phenotypic effects due to a single mutation. (A) Two potential neighborhoods of a green luciferase allele. Amino acids are arranged alphabetically by their full name. The amino-acid states of the ancestral luciferase allele are outlined in boxes, and their identity and position in the sequence are shown below. All amino acid substitutions at the positions that are known to be variable in nature (Stolz et al. 2003) are shown. For simplicity we assume that none of the mutations result in non-viable, “no color” phenotypes. The distribution of phenotypic changes caused by all single amino-acid substitutions of the ancestral allele ***0*** is denoted as ∂***S*_*0*_**. In the left panel, the orange color is accessible. A change to this luciferase for any reason is expected to shift its emission toward orange. In the right panel, most mutations move the phenotype toward blue. A shift toward orange would therefore not be expected by chance or by indirect selection on a trait that is *not* strongly correlated with the color trait under consideration. (B) Comparing the effect of an observed mutation with a neighborhood of possible mutations. White and black boxes indicate the ancestral and derived (mutant) alleles, respectively; their corresponding phenotypes are on the Y-axis. Grey distributions show frequencies of phenotypic changes estimated from each mutational neighborhood. These distributions can be used to estimate the probabilities that a random mutation of the ancestral allele will move the phenotype toward or away from the derived state for this particular trait.

### Multiple-mutation scenario

Now let us consider a more common case. Typically, we have a set of several substitutions that cause a phenotypic difference between the ancestral and derived phenotypes (the positively-selected mutations of the luciferase protein in this example). We have inferred (or know) the ancestral state, but we do not know the temporal order of these substitutions, or the evolutionary paths taken by the population. In particular, we do not know whether the number of substitutions observed between the ancestral and derived sequences is typical, given the amount of evolutionary time that elapsed in the course of the observed shift from green to orange. Given that the actual path taken from the ancestral to the derived allele is difficult or impossible to reconstruct, we propose the following extension of the single-mutation accessibility test. Consider phenotypic measurements for a random subset of genotypes separated by the same number of mutations from the ancestral genotype as the derived genotype. For simplicity, only mutations at the sites found to be different between the ancestral and derived alleles will be taken into account; mutations at all other sites are assumed not to change the phenotype (this assumption can be tested experimentally, and if it proves incorrect, mutations at additional sites can be considered as well). These measurements of phenotypic states will be used to estimate a phenotype probability distribution, which in turn enables a p-value assignment to the derived state’s phenotype in the absence of direct selection.

This type of reasoning, based on collecting information about likely and unlikely changes in the value of the observed trait, can also be extended to the situations when information about evolutionary paths is available. This may be the case in laboratory evolution experiments, where evolving populations can be monitored at regular intervals (Szendro et al. 2013; Jiang et al. 2013; González et al. 2015). In this scenario, the likelihood of each phenotypic change along the path (a member of the ensemble of paths connecting the ancestral and derived alleles) can be assessed using probability distributions constructed from observations of phenotypic states of mutational neighborhoods for each state along the observed path (Fig. 5). Even a single low-likelihood move anywhere along the evolutionary path will constitute strong evidence for the direct selection scenario on the trait under observation. Furthermore, weaker evidence from several steps along the path can be combined by considering their joint probability, which should increase the overall statistical power of the null (indirect selection only) hypothesis test.

**Figure 5.**
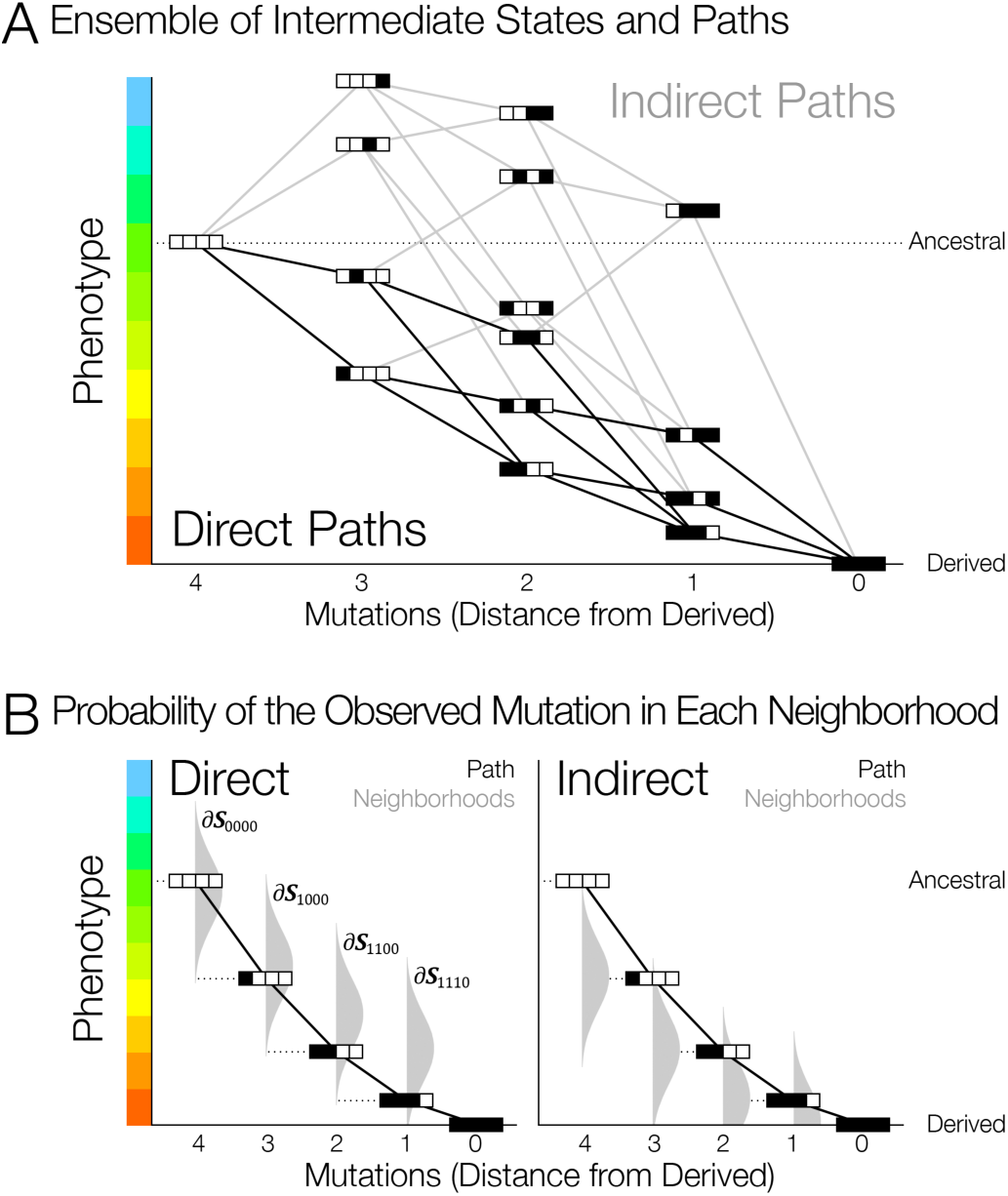
Using mutational accessibility along evolutionary paths to infer the phenotypic target of selection. (A) The ensemble of paths showing all possible intermediate states connecting the ancestral allele and the derived allele. The value of the trait changes monotonically among some of the paths, while on others there is at least one reversal of the trait - a potential indication of indirect rather than direct selection if this type of path is commonly observed. Note that here we assume for simplicity that there are no strongly deleterious mutations, which would not produce a phenotype. (B) Estimating the probability of making the observed mutational substitution at each step along the observed path. For each intermediate allele, we calculate the probability of making the observed substitution using the current allele’s mutational neighborhood. In the left panel, all successive substitutions are unlikely, making direct selection the most probable scenario. In contrast, in the right panel each mutation is likely (according to the distribution of phenotypic changes in the mutational neighborhoods), meaning that the accessibility test gives no reason to reject the indirect selection scenario.

## 4. Challenges facing molecular tests of adaptation

### Phenotypic measurements

Any test of phenotypic evolution rests on our ability to measure phenotypic effects accurately. Phenotypes measured in the lab must resemble their natural counterparts and must be independent of historical, biological, or experimental context in ways that obscure their natural effects.

### Size of phenotype space

The approach we have described is only useful for genetically simple traits, whose variation depends on a small number of genomic loci (i.e., mutations elsewhere do not change the phenotype under study). Many natural adaptations have simple genetic underpinnings, and many examples of molecular, regulatory, and network evolution can be studied mutation-by-mutation (Stern and Orgogozo 2008; Bell 2009; Conte et al. 2012). Indeed it is simple traits that benefit most from explicit genetic analysis, as their evolution may be constrained by their genetics. In these traits, evolutionary processes can be subject to explicit molecular investigations of phenotype space provided one can devise appropriate large-scale phenotype assays.

### Reconstruction of ancestral states and paths

Accurate inference of ancestral states and corresponding evolutionary trajectories is a serious challenge for molecular tests of adaptation. Outside of lineages whose evolution can be observed in real time, the ancestral sequence and the temporal order of mutations could be difficult or impossible to reconstruct (Maynard Smith and Haigh 1974; Gillespie 2000). Ancestral alleles can sometimes remain in a population, but we will only find these cases circumstantially; they cannot underpin a general approach (Springer and Crespi 2007; Rebeiz et al. 2009). Tests of direct and indirect selection in molecular adaptation will therefore have to allow for multiple ancestral sequence candidates and multiple evolutionary paths.

### Population genetics assumptions

Our approach is based on the SSWM assumption – mutations arise one at a time and either fix or disappear, keeping the population monomorphic at all times. Depending on the strength of selection, either the first beneficial mutation discovered will always fix or, somewhat more realistically, the fixation probabilities of beneficial mutations will be proportional to their selection coefficients (Gillespie 1984). In reality, populations may be polymorphic, which could significantly complicate the analysis (although a certain degree of polymorphism can be tolerated by considering the most common sequence variants found in the population). Furthermore, in small populations substitutions may occur due to genetic drift rather than selective advantage, although standard tests to distinguish selection from neutrality (such as McDonald-Kreitman and dN/dS) should help address this problem. Finally, the effects of fluctuating selection and, more generally, past environmental changes may be difficult to capture in artificial conditions.

## 5. Conclusions: Testing adaptation itself

When we call a trait an adaptation, we imagine selection on a particular aspect of phenotypic variation (“hummingbird-pollinated flowers are red because…” or “cave fish lose their eyes because…”). Adaptive explanations invoke specific historical scenarios – both a particular form of selection and, as importantly, assumptions about evolutionary constraints or lack thereof. Adaptation is the center of biology, and determining which phenotypes were direct targets of selection is the key to testing adaptation. We cannot verify an adaptive hypothesis without eliminating the alternative: evolution as a response to selection on a correlated trait (Williams 1966; Gould and Lewontin 1979; Nielsen 2009). A growing number of studies are using artificial mutagenesis to uncover the range of functional phenotypic variation available by mutation. These methods could in principle also be used to verify direct selection, if we could identify unambiguous hallmarks of adaptation. The evolution of a phenotype away from mutationally accessible states is a feature of adaptation by direct selection that could potentially be estimated. Measuring mutational accessibility is not trivial, but can be achieved using existing molecular biology tools. Conveniently, the only traits that truly need genotype-phenotype analysis are genetically the simplest and most tractable (Grafen 1984; Springer et al. 2011). Thus molecular genetics can overcome the problems posed by pleiotropy and phenotypic correlations, and satisfy the deepest goal of evolutionary research – verifying that a phenotypic difference was a direct target of selection and can therefore be considered an adaptation.

## Acknowledgements

SAS is grateful to WJ Swanson, J Tyerman, F Breden and HD Bradshaw for helpful discussions. An NSERC PGSD International Graduate Fellowship to SAS supported this work. MM was supported by NIH grant F32 GM116217. AVM and SAS are grateful to Pierre Pontarotti for his extraordinary hospitality, which enabled the exchange of ideas that ultimately made this work possible.

